# Fulfilling the promise of Mendelian randomization

**DOI:** 10.1101/018150

**Authors:** Joseph K. Pickrell

## 1 Introduction

Many important questions in medicine involve questions about *causality*. For example, do low levels of high-density lipoproteins (HDL) cause heart disease? Does high body mass index (BMI) cause type 2 diabetes? Or are these traits simply correlated in the population for other reasons? The gold standard study design for answering this type of question is the randomized controlled trial, which requires identifying an intervention that influences the potential causal factor (for example a drug), randomly assigning individuals to receive the intervention, and monitoring outcomes. In this study design, the use of randomization allows one to isolate the causal effect of a single variable (e.g. the drug) from the effects of potential confounding variables. However, randomized controlled trials are sometimes impractical (for example, no intervention may exist), expensive, or indeed unethical (for a tongue-in-cheek example, see Smith and Pell [31]).

In the absence of a randomized trial, some hints about the causal relationship between traits can come from observational epidemiology. If two variables are correlated in the population (e.g. low HDL cholesterol is correlated with higher risk of heart disease [14]) then this might indicate a relationship worth examining further. On the other hand, observational correlations are potentially confounded by a large number of factors (e.g. Taubes [34] and references therein), and following up non-causal relationships with randomized controlled trials leads to large investments of time and money with little payoff. Classic examples of this problem include trials of hormone replacement therapy in cardiovascular disease [29] or beta-carotene in lung cancer [36].

A useful attack on the problem of confounding in observational epidemiology comes from genetics, and is now called “Mendelian randomization” (the method is generally attributed to Katan [19], but see Katan [20]). The intuition is as follows: if two traits are causally related, a genetic variant that influences the first trait (for example, BMI) is expected to have a “knock-on” effect on the second trait (for example, type 2 diabetes), while this is not the case if the two traits are correlated for other reasons. Since genetic variants are inherited at conception and (to a first approximation) do not change over the course of an individual’s lifetime, the random inheritance of a genetic variant acts like the randomization in a clinical trial to help avoid the effects of confounding. In the simplest form, Mendelian randomization thus simply involves testing to see if a genetic variant has effects on two traits [19]. More elaborate methods allow one to estimate the magnitude of the causal effect of one trait on the other, and to include multiple genetic variants; for more in-depth description of the statistical methodology, see Didelez and Sheehan [11], VanderWeele et al. [39] and Davey Smith and Hemani [9].

The Mendelian randomization approach is now extremely popular (e.g. [4, 5, 10, 13, 16, 17, 22, 24, 27, 41, 42]). For example, this approach has recently been used to argue that alcohol consumption increases risk of heart disease [16], and that increasing serum iron levels decreases risk of Parkinson’s disease [27]. These findings have potentially important public health implications.

However, it is well-known that causal inference using Mendelian randomization requires strong assumptions about the physiological mechanisms by which genetic variants exert their effects ([9, 39]. Specifically, if a genetic variant influences two phenotypes through different mechanisms, then Mendelian randomization will give misleading results. For example, a genetic variant in the gene LGR4 influences both bone mineral density and risk of gallbladder cancer [33], but it seems implausible to suggest that low bone mineral density causes cancer. Instead, it seems more likely that this genetic variant alters some molecular pathway (or multiple pathways) that is important for both phenotypes. Here, we describe two pieces of evidence that make us moderately skeptical of most of the causal claims made to date using Mendelian randomization, and suggest some ways forward.

## 2 Two reasons for skepticism

The first reason for skepticism is that in the nearly 30 years of Mendelian randomization, arguably no new causal relationship has been identified with this approach and subsequently verified in a randomized controlled trial. We assembled a list of 29 applications of Mendelian randomization from two recent reviews [3, 9] (Supplementary Table 1). Of these 29 pairs of traits, we identified five that were also targets of randomized controlled trials. In nearly all of these case, the Mendelian randomization study was performed *after* the randomized controlled trial, and generally confirmed the results of the trial (Table 1). For example, Mendelian randomization studies have provided additional evidence that BMI causally influences atherosclerosis [21], and that HDL levels do not appear to causally influence risk of cardiovascular disease [41]. These studies have provided important, independent evidence about the relationships between these traits, but it seems the most rigorous test of the Mendelian randomization methodology would be to identify a new causal relationship and then to subsequently confirm it with a controlled trial.

**Table 1.** List of causal relationships evaluated by Mendelian randomization and randomized controlled trials, compiled from Table 1 of Davey Smith and Hemani [9] and Table 1 of Burgess and Thompson [3]. For the complete list of studies, see Supplementary Table 1. MR = Mendelian randomization, RCT = randomized controlled trial.

| Trait | Outcome | MR effect | Reference | RCT effect | Reference | Performed first |
| --- | --- | --- | --- | --- | --- | --- |
| cholesterol levels | cancer | None | [38] | None | [7] | RCT |
| HDL cholesterol | myocardial infarction | None | [41] | None | [1] | RCT |
| homocysteine | stroke | + | [5] | Conflicting | [18, 32] | MR |
| BMI | carotid thickness | + | [21] | + | [15, 30] | RCT |
| interuterine folate | neural tube defects | - | [8] | - | [25] | RCT |

The one case where Mendelian randomization provided support for a causal relationship prior to the publication of a randomized controlled trial was the case of homocysteine levels and stroke [5]. This study inferred that increased homocysteine levels cause higher stroke risk. However, randomized controlled trials of folic acid (which reduces levels of circulating homocysteine) have produced conflicting results [18, 32]. There are a number of potential explanations for this (indeed, the robustness of the Mendelian randomization study has been questioned [23]), but overall the track record of Mendelian randomization is sparse.

The second reason for skepticism is more fundamental. Specifically, one of the core assumptions when using Mendelian randomization to infer causality is that the genetic variants used in the study have a direct effect on one trait (the “causal” trait), but only an indirect effect on the other (the “caused” trait). That is, the genetic variants used in the study are assumed to have no influence on confounding factors that influences both traits. This is often referred to as an assumption of no “pleiotropy” [9]. However, it is difficult to know *a priori* whether this assumption is valid, and recent work in human genetics has shown that genetic variants often have effects on different traits. For example, genetic variants that influence one autoimmune disease also often influence others [6], and variants that influence one lipid trait also often influence other lipid traits [35]. This is not limited to traits that are intuitively related, however–large genome-wide association studies are finding that genetic variants often influence many aspects of physiology [2]. For example, genetic variants that influence HDL cholesterol levels have correlated effects on whether an individual went to college [2]; a naive interpretation of this might suggest the (rather nonsensical) conclusion that HDL-raising drugs should increase education levels.

It is sometimes suggested that using a large number of genetic variants in Mendelian randomization (combined into a single score) offers a way to partially avoid this problem (e.g. [9, 17]). To test this, we performed simulations of such an approach in a situation where two traits are not in fact causally related (Figure 1A, see Supplementary Information for details). We found that even a small number of pleiotropic loci (those that influence a confounding variable) leads to false positives, and that this problem grows worse as sample sizes increase (Figure 1B). In empirical data, this effect has been seen by Evans et al. [12], who noted that a related approach identified several apparently spurious causal relationships between traits. Before making a strong causal conclusion from a Mendelian randomization study, then, it seems necessary to have a good molecular and physiological understanding of the effects of all the genetic variants used in the study. For many traits of interest, our understanding of biology is not strong enough for this to be reasonable.

**Figure 1.**
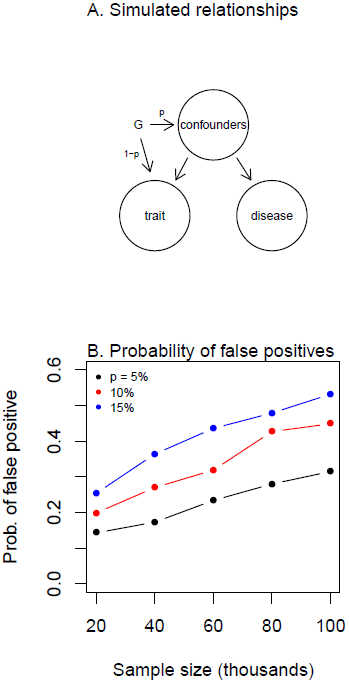
Simulations of Mendelian randomization in the absence of a causal relationship between a trait and a disease. **A. Simulated scenario.** For each simulation, we simulated 100 genetic variants (G), which either 1) influence a trait or 2) influence both the trait and the disease, with proportions 1 − *p* and *p*, respectively. Arrows represent causal relationships [26]. The confounders are not directly simulated; see Supplementary Material for details. **B. Probability of incorrectly inferring a causal relationship between a trait and a disease in this situation**. We simulated Mendelian randomization studies with different sample sizes and values of *p*, and in each simulation tested for a causal relationship between the trait and the disease. Each point shows the proportion of 1,000 independent simulations where the causal effect of the trait on the disease was “significant” as judged by *P* < 0.05. In all simulations, we assumed that the effects of a variant on the trait and on the disease are uncorrelated; see Figure S1 for simulations where these effect sizes are correlated.

## 3 Fulfilling the promise of Mendelian randomization

The principle of Mendelian randomization, however, remains tremendously promising. What are the ways forward? One possibility is to treat negative results from Mendelian randomization with more confidence than positive results–if two traits are correlated in the population but a Mendelian randomization study suggests they are not causally related, this is evidence that the population-level correlation is driven by some unobserved confounder [39]. Another approach might be to focus on developing a detailed molecular understanding of variants in an individual gene and their downstream physiological effects; this approach is particularly useful if the gene is a potential drug target [28]. Finally, another possibility comes from the ongoing development of new statistical methods like bi-directional Mendelian randomization (e.g. [37, 40]), which use many genetic variants to increase the robustness of causal inference. These methods are currently in early stages of development and evaluation. Until they reach maturity, reports of causal relationships identified through Mendelian randomization should be treated with the same skepticism as such claims made from observational epidemiology.

## 4 Acknowledgements.

We thank Molly Przeworski for helpful comments on an earlier draft of this manuscript.

